# Space matters: virtual pedestrians with mobility constraints affect individuals’ avoidance behaviours

**DOI:** 10.1101/2024.10.21.619458

**Authors:** Mohammadamin Nikmanesh, Michael E. Cinelli, Daniel S. Marigold

**Affiliations:** Biomedical Physiology and Kinesiology, Simon Fraser University, Burnaby, BC V5A 1S6, Canada; Institute for Neuroscience and Neurotechnology, Simon Fraser University, Burnaby, BC V5A 1S6, Canada; Department of Kinesiology and Physical Education, Wilfrid Laurier University, Waterloo, ON, Canada

## Abstract

Walking in urban settings requires people to negotiate crowds. In these situations, people typically want to maintain a level of personal space around themselves. Recent work on one-versus-one interactions demonstrated that whether one of the pedestrians looked distracted or interacted with an object (e.g., stroller, bike) predicted the medial-lateral separation between them as they walked past each other. However, this work did not distinguish between the type of object interaction (or mobility constraint) and thus, it is unclear whether different constraints have different effects on avoidance behaviours. Here we tested the hypothesis that the type of object an approaching pedestrian held or pushed would affect the extent of path deviation, which would also depend on whether that pedestrian was distracted. To address this hypothesis, we created an immersive virtual environment that consisted of a 3.5-m-wide paved urban path. Participants had to walk and avoid colliding with approaching virtual pedestrians that often held a shopping bag or pushed a bike or stroller while looking straight ahead or off to the side as if distracted. Distraction did not affect avoidance behaviours. However, participants increased medial-lateral separation with the virtual pedestrian at the time of crossing when a stroller was present compared to the other mobility constraints. The type of mobility constraint also differentially affected onset of deviation and rate of progression before and after a path deviation. These results support the idea that characteristics of the obstacle to avoid (in this case, a virtual pedestrian) influence collision avoidance behaviours.

## INTRODUCTION

In urban areas, people must often adjust their walking pattern to avoid collisions with obstacles and surrounding pedestrians. A wide variety of factors have the potential to influence these adjustments, including age (Rapos et al. 2021), body size (Bourgaize et al. 2021), heading direction or approach angle (Huber et al. 2014), and crowd density (Bruneau et al. 2015; Koilias et al. 2020). Understanding what characteristics impact avoidance behaviours is important for developing and refining computational models used to predict or explain pedestrian walking trajectories, as well as models of crowd simulation.

People typically maintain a certain level of personal space when interacting with others, whether standing and having a conversation (Hecht et al. 2019) or walking (Gérin-Lajoie et al. 2005). Defined as the immediate area surrounding an individual, personal space is an instinctive and dynamic protective buffer that pedestrians maintain to prevent collisions (Bourgaize et al. 2021; Gérin-Lajoie et al. 2005; Gorrini et al. 2014; Liu et al. 2019). Personal space is a margin of safety also observed in animals, such as monkeys, where uninvited intrusions into this zone evoke defensive behaviours (Graziano and Cooke 2006). Walking pedestrians tend to maintain an elliptical personal space envelope, with that space extending ∼2 m in front and ∼0.5 m to one side (Gérin-Lajoie et al. 2005); the shape of this space changes to circular with a radius of ∼1 m when people are stopped and having a conversation (Hecht et al. 2019). In higher-density settings, where personal space violations are more frequent, pedestrians increase walking speed and are more likely to change walking direction, possibly to escape the uncomfortable (or stressful) situation (Frohnwieser et al. 2013). In addition, people demonstrate greater medial-lateral (ML) personal space when walking past a larger-, compared to smaller-sized, stationary person in the laboratory (Bourgaize et al. 2021); ML personal space shrinks when the stationary person faces to the left or right rather than forward or backward, implying that people consider the most lateral edge of an interferer when altering their walking trajectory. Taken together, one could argue that ensuring some degree of personal space is a primary objective while navigating environments populated with other pedestrians.

Recently, we investigated unscripted pedestrian walking behaviours on a busy urban path (Nikmanesh et al. 2024). We used deep learning algorithms to identify and extract pedestrian walking trajectories from videos of the urban path where two pedestrians approached each other from opposite ends. We found that the ML distance between the approaching pedestrians and surrounding crowd size predicted the likelihood of a path deviation. Our previous findings also revealed that when one of the pedestrians looked distracted, the ML separation between them was larger as they walked past each other (Nikmanesh et al. 2024). Laboratory-based experiments have repeatedly demonstrated that distraction affects walking behaviours (Gérin-Lajoie et al. 2006; Lamberg and Muratori 2012; Lin and Huang 2017; Murakami et al. 2022; Souza Silva et al. 2019, 2020). Distracted walkers are more likely to increase their lateral deviation (Lamberg and Muratori 2012), and the non-distracted walker may increase how much they deviate to avoid them (Murakami et al. 2022). Walkers distracted with a cell phone are slower to detect approaching virtual pedestrians and are more likely to collide with them (Souza Silva et al. 2019, 2020). Interestingly, we also found that when one of the pedestrians interacted with an object (e.g., stroller, bike) or animal, the ML separation between them as they walked past each other was smaller than with no constraint. It is possible that when one’s movement is constrained, an approaching pedestrian perceives that they are less mobile and thus, can get closer without a risk of collision. Based on our previous study observing real-world interactions, it appears that mobility constraints affect collision avoidance behaviours. However, we did not observe enough instances to distinguish between the type of object interaction (or mobility constraint) and thus, it is unclear whether different constraints have different effects on these behaviours.

In the present study, we developed a virtual reality (VR) environment that replicated the physical characteristics of the real-world location—in Vancouver, British Columbia—used in our previous research study (Nikmanesh et al. 2024). In the current virtual environment, we had a virtual pedestrian approach the participants while holding a shopping bag or pushing a bike or stroller and either look straight ahead or off to the side as if distracted. We tested the hypothesis that the type of constraint the approaching pedestrian held or pushed would affect the extent of path deviation (i.e., ML separation at the time of crossing), which would also depend on whether that pedestrian was distracted. We also determined how these mobility constraints and distraction affected different factors that could contribute to the extent of deviation (e.g., onset of deviation, progression rate).

## METHODS

### Participants

Fifteen young adults (five males, 10 females; mean ± SD age = 28.0 ± 4.0 years, height = 162.9 ± 9.1 cm) participated in the experiment. Participants did not have any known visual, neurological, muscular, or joint disorder that could affect their walking behaviour, and did not regularly suffer from motion sickness or have a history of virtual reality intolerance. The Office of Research Ethics at Simon Fraser University approved the study protocol, and participants provided informed, written consent prior to data collection.

### Experimental set-up and VR environment

We conducted the experiment in a large room with a 7 m long, 4 m wide pathway cleared along the middle. Participants had to walk in a virtual environment and avoid colliding with an approaching virtual pedestrian. We created the virtual environment using Unreal Engine 5.4 and presented it to participants outfitted with the HTC Vive Focus 3 VR head-mounted display, which was free from wires and allowed participants to have unimpeded movement. The VR environment featured a 3.5-m-wide paved urban path, with benches and a stone wall on one side and grass, sand, wooden posts, and logs on the other (see Figure 1). The setting resembled the location of our past work in Vancouver, British Columbia, including similar path width and placement of objects. A square at the beginning of the walking path denoted the start location. The VR system enabled us to record (at a dynamic sampling frequency between 80 and 90 Hz to minimize lag) and extract kinematic data from each participant and the virtual pedestrian within the environment. Specifically, it estimated the location of each participant’s head in space, which was necessary to calculate our dependent variables.

**Figure 1.**
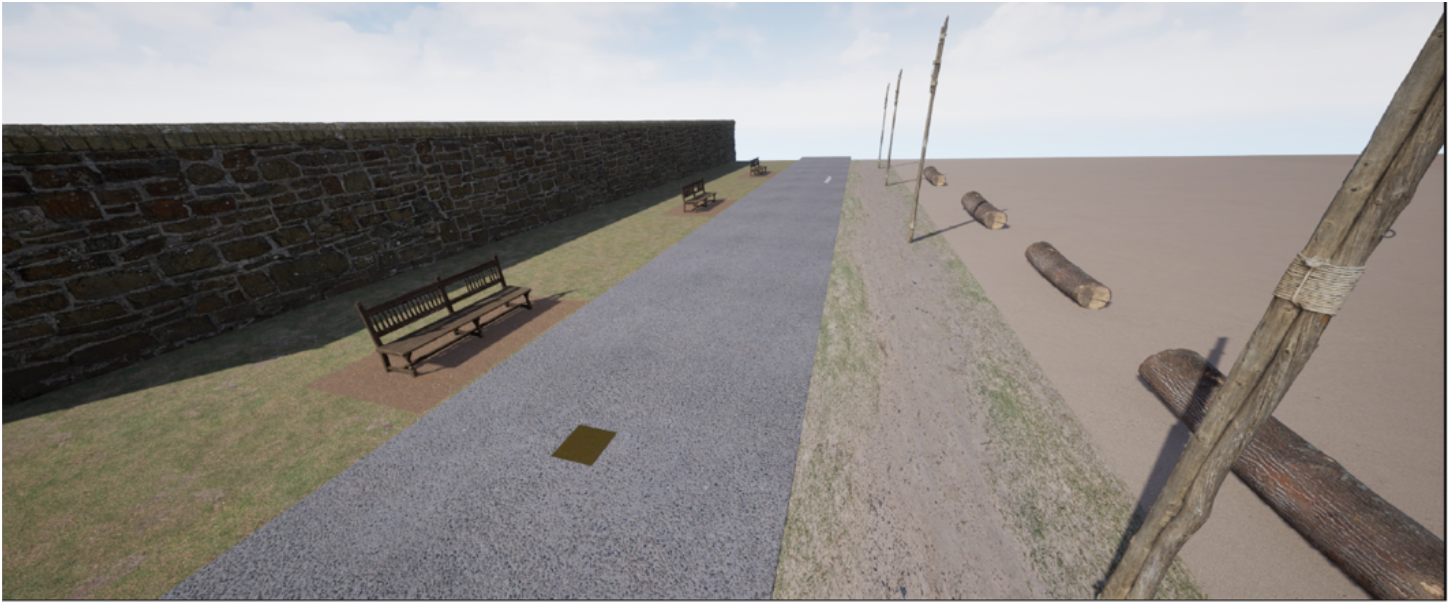
Illustration of the VR Environment.

Participants encountered one of four pedestrian mobility constraint conditions (see Figure 2): pedestrian walking without any constraint (control condition), pedestrian walking while holding a shopping bag, pedestrian pushing a bicycle, and pedestrian pushing a stroller. The virtual pedestrian either walked with their head rotated such that they looked away from the path and participant (i.e., distracted condition) or maintained their head (and gaze) facing forward along the path (i.e., not distracted condition). The virtual pedestrian exhibited random fluctuations of ±1 cm in the ML direction during movement to simulate natural variability in walking patterns.

**Figure 2.**
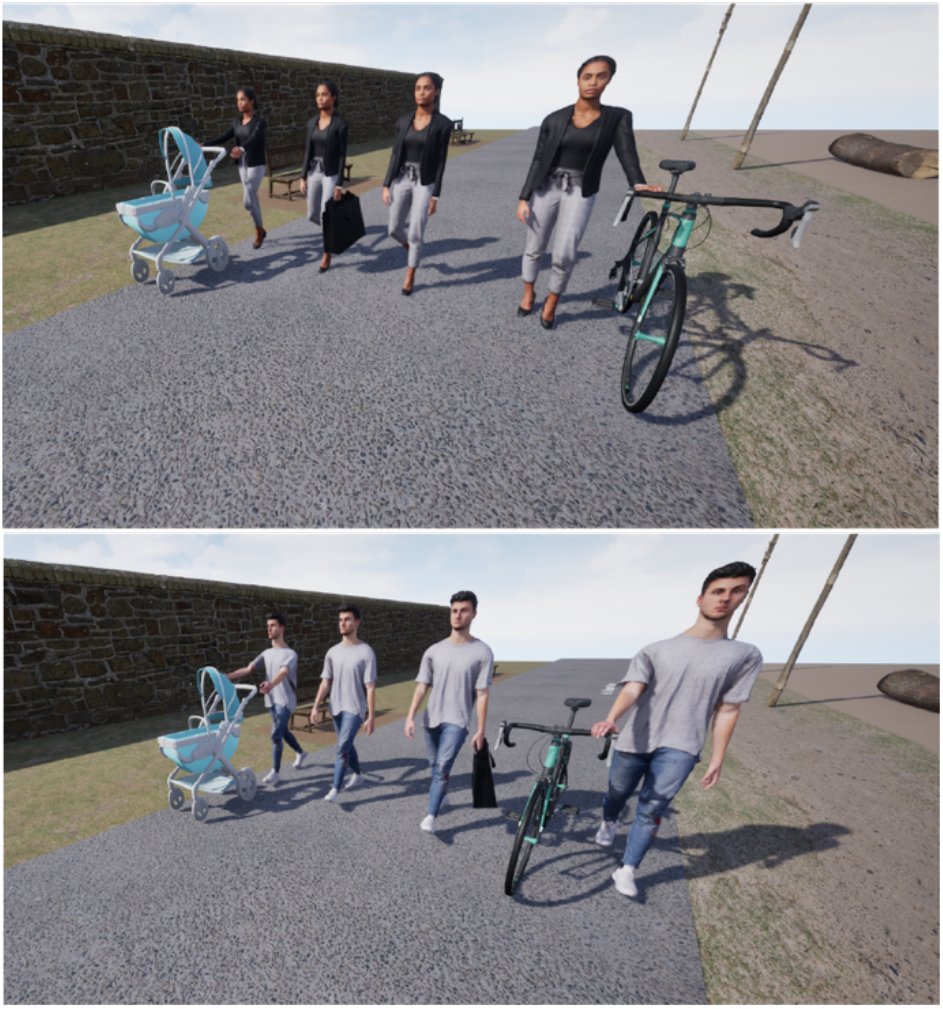
Virtual pedestrians and mobility constraints. Top: Female virtual pedestrian (from left to right: pushing a stroller, not interacting with an object, carrying a shopping bag, and pushing a bicycle). Bottom: Male virtual pedestrian (from left to right: pushing a stroller, not interacting with an object, carrying a shopping bag, and pushing a bicycle).

### Procedure

Prior to the start of the experiment, we asked participants to select one of two virtual pedestrians that they would like to interact with during the task, one resembling a female and the other a male. The selected pedestrian remained consistent throughout the experiment. For the experiment, we instructed participants to walk past the pedestrian without colliding with it but otherwise did not specify how to walk.

Participants started each walking trial by standing on a virtual square. To maintain participant’s focus on the task, the square’s position alternated between two locations that were ± 40 cm from the centre line of the walking path. After the participant walked 0.5 m, the virtual pedestrian appeared 8.5 m ahead and approached at a constant speed of between 1.0 and 1.2 m/s (randomly chosen each trial). Given the two different start locations, the virtual pedestrian began either on the left or the right of the participant. The virtual pedestrian disappeared after the participant walked past and then the participant returned to a starting square. Participants received a 10-second break between walking trials.

Participants completed 64 walking trials. This included 12 trials for each pedestrian mobility constraint condition. Additionally, we included 16 catch trials, in which the virtual pedestrian changed their locomotor trajectory and walked toward the participant. The catch trials ensured that participants did not assume the virtual pedestrian was unresponsive or always walked along a straight path. Half of the trials simulated the virtual pedestrian being distracted (i.e., looked away), while in the other half, the pedestrian maintained forward gaze. We presented trials in four blocks of 16. A block consisted of four catch trials and three trials of each of the four mobility constraint conditions, all randomized. After each block of trials, an experimenter asked participants if they had any symptoms of motion sickness, like nausea, headaches, or dizziness. No one reported any of these symptoms.

#### Data analyses

We excluded catch trials and trials where there was a problem (e.g., virtual pedestrian did not appear) prior to our analyses. Subsequently, we filtered the kinematic data with a 4^th^-order, 6 Hz low-pass, zero-lag Butterworth algorithm.

As a primary measure, we calculated the ML separation at the point in time when the participant walked past the virtual pedestrian (i.e., ML separation at time of crossing). ML separation represented the absolute difference between the ML position of the participant (derived from the VR headset and was representative of the centre of the participant) and the virtual pedestrian (the centre point) at the time of crossing. We included all trials in this analysis regardless of whether the participant deviated in their walking trajectory.

We also considered several secondary measures that could contribute to avoidance behaviour. As an initial step, we determined if a participant deviated to avoid the virtual pedestrian. To accomplish this, we calculated the first derivative of the participant’s ML position to detect local minima, where the ML position reached a low point and started to increase again. These local minima, plus the participant’s initial position were considered as potential deviation points. We did not consider points within 0.5 s before the crossing point because there would not be enough time for the participant to initiate a deviation. For each potential deviation point, we set a deviation threshold, the value of which represented the ML position of this point plus 2 standard deviations of the ML position calculated 0.5 s before the virtual pedestrian appeared. We classified a trial as a deviation if the absolute ML position of the best candidate point had a lower ML value compared to the crossing point, and no subsequent point had a smaller ML value. If the ML position of a new candidate point was below the calculated threshold of the previous point, and no better candidates were identified after that, this point was classified as the onset of deviation (see Figure 3). If these conditions were not met, we considered the trial as having no deviation. The corresponding time of deviation was recorded as the time from the start of the trial to this point. This approach allowed us to robustly identify the best candidate for deviation while accounting for noise and ensuring accuracy before crossing occurred.

**Figure 3.**
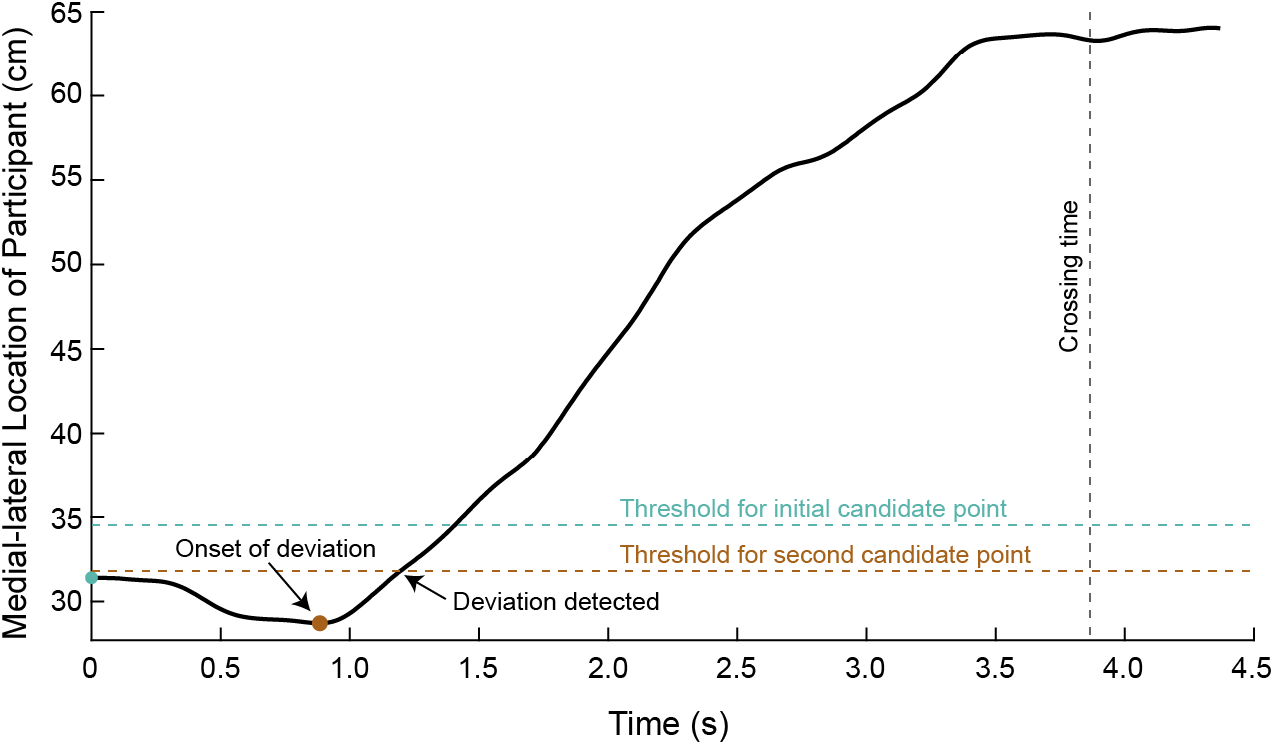
Illustration of the approach to determine the presence and onset of a deviation in walking trajectory. For each trial, we identified the initial starting position and local minima based on the derivative of the medial-lateral position of the participant as candidate points. For each point, we set a deviation threshold, which represented the medial-lateral position of this point plus 2 standard deviations of the medial-lateral position calculated 0.5 s before the virtual pedestrian appeared. See text for additional details.

For deviation trials, we calculated:

1. Average rate of deviation, defined as the average instantaneous ML rate (speed in ML direction or difference in ML position over difference in time) between the onset of deviation and the time of crossing.
2. Rate of progression (akin to walking speed) based on two intervals: start of walking to the onset of deviation (i.e., before deviation) and onset of deviation to the time of crossing (i.e., after deviation). Rate of progression involved first determining the instantaneous derivative of the Euclidean distance (anterior-posterior and ML) over time and then taking the average for each of the two intervals.

#### Statistical analyses

We compared the following dependent variables across mobility constraint conditions (control, shopping bag, bike, stroller) and distraction conditions (forward gaze and looking away): average ML separation at the time of crossing, onset of deviation, rate of deviation, and rate of progression (both before and after the onset of deviation). Specifically, we performed separate two-way (Mobility Constraint x Distraction) linear mixed-effects models (with participant as a random effect). We used JMP 16 software (SAS Institute Inc., Cary, NC) with an alpha level of 0.05 for all statistical analyses, and report effect sizes using partial eta squared 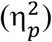. When we found significant mobility constraint effects from the linear mixed-effects models, we performed Tukey’s post hoc tests.

## RESULTS

### The type of object constraining the virtual pedestrian affects ML separation at time of crossing

We compared ML separation between the participant and virtual pedestrian at the time of crossing across four different mobility constraint conditions (control, bike, shopping bag, and stroller) and two distraction conditions (no distraction/forward gaze, and distraction/looking away). The results showed a significant main effect of mobility constraint (F_3, 98_ = 10.95, p = 2.9e-6, 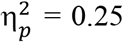), indicating that ML separation at crossing differed depending on the type of mobility constraint (Figure 4). Post hoc comparisons revealed significantly greater ML separation at crossing in the stroller condition compared to all other conditions (p < 0.05). Additionally, ML separation at crossing did not significantly differ between the control, bike, and shopping conditions (p > 0.05). We did not detect a significant main effect of distraction (F_1, 98_ = 0.29, p = 0.593, 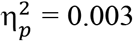) or interaction (F_3, 98_ = 0.80, p = 0.497, 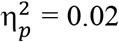). When collapsed across distraction condition, the mean ± SD of the ML separation at crossing for the control, bike, shopping bag, and stroller conditions were 100.9 ± 14.0 cm, 101.4 ± 15.1 cm, 100.0 ± 13.3 cm, and 106.4 ± 14.3 cm, respectively.

**Figure 4.**
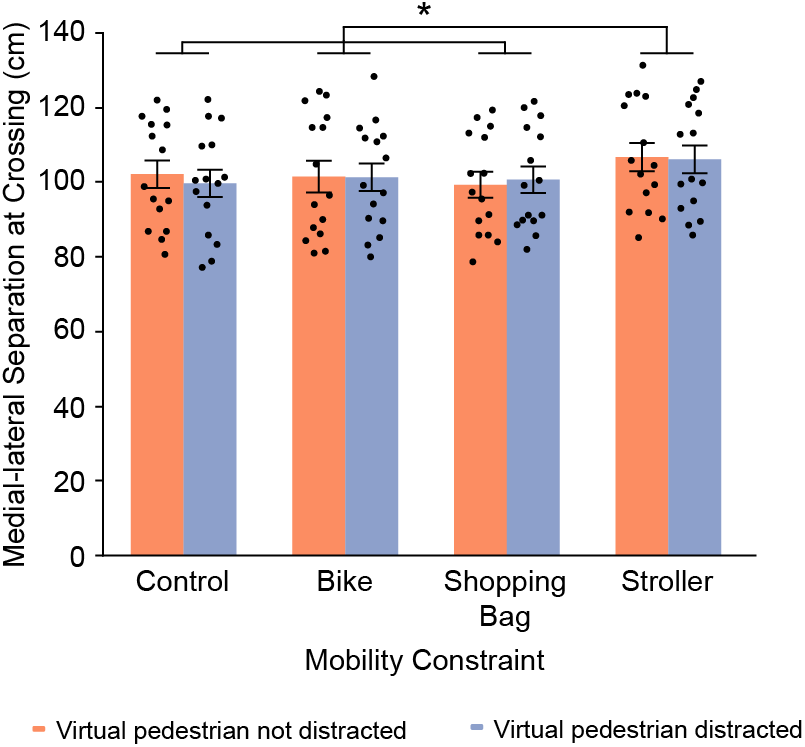
Medial-lateral separation at time of crossing. Group means ± SE of medial-lateral separation at time of crossing across mobility constraint and distraction conditions are shown. Data from each individual participant are superimposed. * Indicates conditions that are significantly different from each other based on post hoc tests related to the main effect of mobility constraint (p < 0.05).

### Onset of deviation depends on the type of constraint on the virtual pedestrian

We first identified trials in which the participants deviated while walking. This happened in most trials. In fact, participants deviated on average 86.6% of trials (range: 61.7 - 100%). Subsequently, we compared the onset of deviation across the four different mobility constraint conditions (control, bike, shopping bag, and stroller) and two distraction conditions (no distraction/forward gaze, and distraction/looking away). The results are shown in Figure 5. Specifically, the results demonstrated a significant main effect of mobility constraint (F_3, 98_ = 4.47, p = 0.0055, 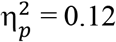), indicating that the onset of deviation varied based on the type of mobility constraint. Post hoc comparisons revealed an earlier onset of deviation in the stroller condition compared to the bike and shopping conditions (p < 0.05). However, onset of deviation in the stroller condition did not differ from the control condition (p > 0.05). Despite the effects of mobility constraint, we did not detect a significant main effect of distraction for onset of deviation (F_1, 98_ = 1.07, p = 0.304, 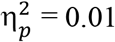). We also did not detect a significant interaction between mobility constraint and distraction (F_3, 98_ = 1.08, p = 0.362, 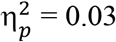).

**Figure 5.**
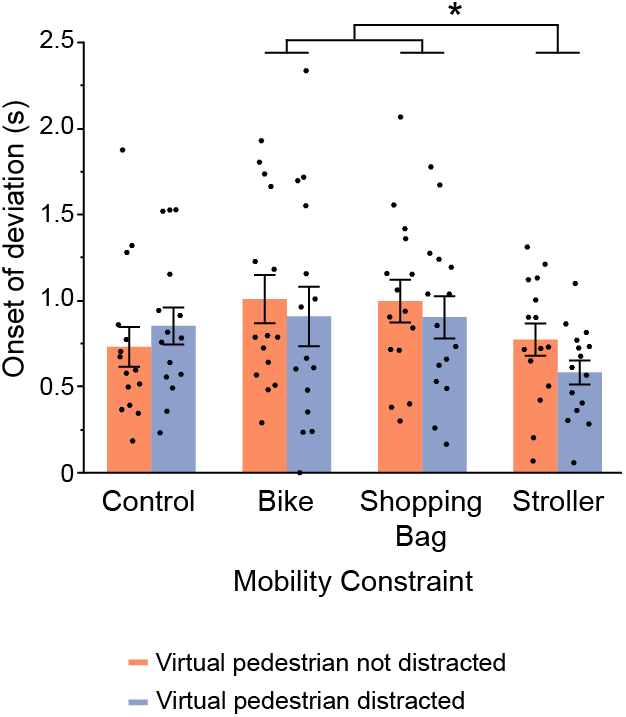
Onset of deviation. Group means ± SE for onset of deviation across mobility constraint and distraction conditions. Data from each individual participant are superimposed. * Indicates conditions that are significantly different from each other based on post hoc tests related to the main effect of mobility constraint (p < 0.05).

### Progression rates depend on the type of virtual pedestrian constraint while walking

Amongst deviation trials, we tested whether mobility constraint and/or distraction affected the rate of deviation or the rate in which participants walked (i.e., progression rate). The results are illustrated in Figure 6. For deviation rate (Figure 6A), we did not detect a significant main effect of mobility constraint (F_3,98_ = 2.32, p = 0.080, 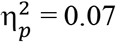) or distraction (F_1,98_ = 0.34, p = 0.559, 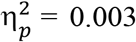), and we also found no significant interaction (F_3,98_ = 0.16, p = 0.926, 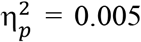). Mobility constraint affected the rate of progression before deviation (Figure 6B; F_3,97_ = 3.24, p = 0.026, 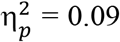) and the rate of progression after deviation (Figure 6C; F_3,98_ = 3.93, p = 0.011, 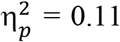). Post hoc comparisons revealed a slower rate of progression before deviation in the stroller condition compared to the control condition and a slower rate of progression after deviation in the bike condition compared to the stroller and control conditions (p < 0.05). However, we did not detect a significant main effect of distraction for progression rate before deviation (F_1,97_ = 0.04, p = 0.838, 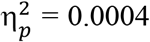) or after deviation (F_1,98_ = 0.24, p = 0.626, 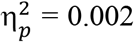). We also did not detect a significant interaction for progression rate before deviation (F_3,97_ = 1.59, p = 0.197, 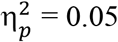) or after deviation (F_3,98_ = 0.53, p = 0.665, 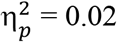).

**Figure 6.**
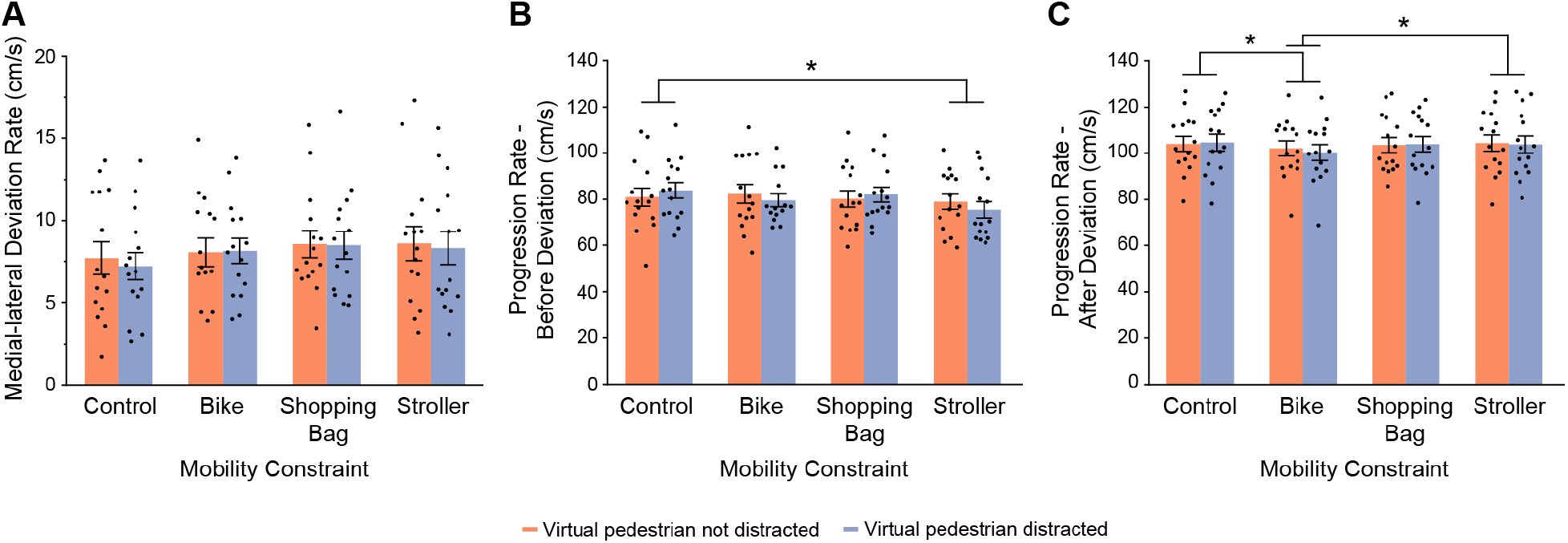
Rates of deviation and forward progression before and after deviation. Group means ± SE for medial-lateral deviation rate (A), progression rate before deviation (B), and progression rate after deviation (C) across mobility constraint and distraction conditions. Data from each individual participant are superimposed. * Indicates conditions that are significantly different from each other based on post hoc tests related to the main effect of mobility constraint (p < 0.05).

We also asked whether progression rate differed before versus after deviation. Since distraction did not affect this measure (as demonstrated in Figure 6B,C), we combined data across distraction conditions and compared progression rates across time (before and after deviation) and mobility constraint conditions. We found that progression rate significantly increased after deviation (103.3 ± 12.8 cm/s) compared to before deviation (80.5 ± 12.5 cm/s), as evident from a significant main effect of time (F_1,98_ = 568.67, p = 1.37e-42, 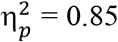). However, we did not detect a significant main effect of mobility constraint (F_3,98_ = 1.54, p = 0.209, 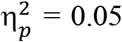) or an interaction (F_3,98_ = 2.55, p = 0.060, 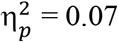).

## DISCUSSION

When walking, people often interact with an object that has the potential to constrain their mobility, like holding a shopping bag or pushing a stroller or bike. In the current study, we used a virtual environment to determine whether the type of mobility constraint, and whether the virtual pedestrian with the constraint appeared distracted, affected collision avoidance behaviours. The mobility constraints used in our experiment defined the characteristics of the virtual pedestrian and consequently, the obstacle that our participants had to avoid. Our results revealed the earliest onset of deviation and the greatest ML separation at time of crossing occurred during the stroller condition. The results also revealed that the stroller condition produced a slower rate of progression prior to deviation and a faster rate of progression following the deviation. Regardless of condition, participants walked faster after deviation compared to before deviation. However, whether the virtual pedestrian appeared distracted or not had no effect on any of our measures. Taken together, our results suggest that characteristics of the obstacle to avoid (in this case, a virtual pedestrian) influence collision avoidance behaviours.

The virtual pedestrians with a mobility constraint affected the ML separation at time of crossing such that participants perceived strollers as requiring more space (Figure 4). It is possible that participants associated a stroller with a (potentially sleeping) child and thus, opted for a wider clearance to avoid disturbing them (despite being in a virtual environment). This would suggest that participants consider affordances of others when controlling behaviours (Gibson 1979). It is also possible that participants considered the bulk of the stroller as being riskier than the shopping bag or condition where the virtual pedestrian did not have a mobility constraint. This implies that obstacle size can affect avoidance behaviours. In support of this idea, people leave a greater ML separation at time of crossing when avoiding a larger interferer compared to a smaller one (Bourgaize et al. 2021; Caplan and Goldman 1981). Furthermore, children demonstrate smaller ML separation at time crossing when interacting with another child or an adult compared to two approaching adults (Rapos et al. 2019); a child’s smaller body size may explain these findings. In addition, pedestrians pushing/pulling luggage can influence the avoidance strategy of the approaching pedestrian without the mobility constraint, such as the extent of path deviation, which depends on the approach angle (Shi et al. 2024). Despite the bike in our study also being bulky, that mobility constraint condition did not evoke the same avoidance strategy. However, the virtual pedestrian always pushed the bike—and likewise, carried the shopping bag—on the side opposite to the approaching pedestrian. This may have led to the perception of it being safer to maintain a smaller ML separation with the virtual pedestrian in these scenarios. The finding that the characteristics of the obstacle (or pedestrian) is important is in line with previous laboratory studies using physical and/or virtual environments. These studies demonstrated that age (Rapos et al. 2019, 2021), body size (Bourgaize et al. 2021; Caplan and Goldman 1981; Hartnett et al. 1974), and whether the obstacle was real versus virtual (Bühler and Lamontagne 2019; Sanz et al. 2015) or was human or inanimate (Sanz et al. 2015; Souza Silva et al. 2018) could influence ML separation at time of crossing. Taken together, the space around a person matters.

The greater ML separation at time of crossing with the stroller contrasts with the direction of the effect observed in our past work on real-world interactions, which demonstrated smaller ML separation at time of crossing when at least one of the approaching pedestrians had a mobility constraint (Nikmanesh et al. 2024). This difference may relate to the environment, where one is in the real-world and the other in a virtual world. Interestingly, studies have found greater ML separation at time of crossing when a person circumvents a static virtual pedestrian compared to a real pedestrian (Bühler and Lamontagne 2019; Sanz et al. 2015). This observation may relate to the fact that people tend to underestimate distances in VR (Kelly et al. 2013; Willemsen et al. 2009) or because they cannot peripherally see their bodies in VR due to the reduced field of view (Hackney et al. 2020). Unfortunately, in our real-world study (Nikmanesh et al. 2024), there were not enough instances of different types of mobility constraints to determine if pedestrians increased ML separation with a stroller but decreased ML separation with other constraints to drive the overall effect.

The presence of a mobility constraint also affected the temporal aspect of avoidance and progression rates differed before and after a path deviation. Specifically, participants initiated their path deviation earlier in the stroller condition compared to the other mobility constraint conditions (Figure 5). This earlier onset of deviation was likely related to the greater ML separation at time of crossing in this condition (Pfaff and Cinelli 2018), as when coupled with a similar rate of ML deviation (Figure 6A), participants required more time to reach their desired ML separation. Participants also increased their rate of progression after relative to before deviation in all conditions. This may suggest that participants felt safer walking at a relatively quicker pace after they were more certain they would avoid a collision.

The virtual pedestrian sometimes appearing distracted did not affect avoidance behaviours, which contrasts with our previous work on real-world pedestrian interactions (Nikmanesh et al. 2024), as well as laboratory-based studies on distraction (Gérin-Lajoie et al. 2006; Lamberg and Muratori 2012; Lin and Huang 2017; Murakami et al. 2022; Souza Silva et al. 2019, 2020). Given these previous results, we had expected participants to exhibit greater ML separation when walking past a virtual pedestrian that looked away from the path (thus appearing distracted) compared to one that looked straight ahead. In this scenario, we reasoned that participants would assume the pedestrian not paying attention to the path ahead would be more likely to have an irregular walking trajectory (Lamberg and Muratori 2012) and/or less likely to contribute to the avoidance behaviour, thus requiring greater deviation to ensure safety. However, it is possible that participants did not perceive the virtual pedestrian’s head rotation. Without capturing gaze, we could not determine where participants looked, and specifically whether they fixated the virtual pedestrian’s head. Regardless, our results emphasize the importance of mobility constraints, rather than attentional factors, in affecting avoidance behaviours.

There are at least two limitations of our experiment. First, our VR environment represented an isolated situation. Although we designed the environment to closely mirror a real-world location, it did not fully capture the complexity of it or real-world environments in general, such as the presence of multiple pedestrians, different terrain, or visual distractions. Second, VR limits any threats to injury if one was unsuccessful at avoiding a collision.

In conclusion, our findings underscore the unique challenges posed by mobility constraints, particularly strollers, in pedestrian environments. Our results revealed that pedestrians naturally adjusted their walking trajectory at different times and to different extents based on the type of an approaching pedestrian’s mobility constraint. This insight could improve computational models of avoidance behaviours and VR simulations. For instance, the Behavioural Dynamics model treats people and obstacles as simply points in space (Fajen and Warren 2003) and thus, does not consider these constraints. In addition, the integration of these considerations into urban planning may enhance the design of more inclusive and safer pedestrian environments. For instance, urban planners could design pedestrian walkways, crosswalks, and other shared spaces with the needs of people using mobility aids or pushing objects (like a stroller) in mind. Our findings indicate that more space should be allocated to accommodate these people, helping to prevent collisions and ensure smoother pedestrian flow in crowded areas. Future research could explore using different (and more complicated) environments and other mobility constraints, like someone carrying a child, holding a cellular phone, or pushing a walker.

## REFERENCES

Bourgaize, S.M., McFadyen, B.J., & Cinelli, M.E. Collision avoidance behaviours when circumventing people of different sizes in various positions and locations. J. Mot. Behav. 53, 166–175 (2021). 10.1080/00222895.2020.1742083

Bühler, M.A., & Lamontagne, A. Locomotor circumvention strategies in response to static pedestrians in a virtual and physical environment. Gait. Posture. 68, 201–206 (2019). 10.1016/j.gaitpost.2018.10.004

Bruneau, J., Olivier, A-H., & Pettré, J. Going through, going around: a study on individual avoidance of groups. IEEE. Trans. Vis. Comput. Graph. 21, 520–528 (2015). 10.1109/TVCG.2015.2391862.

Caplan, M.E., & Goldman, M. Personal space violations as a function of height. J. Soc. Psychol. 114, 167–171 (1981). doi:10.1080/00224545.1981.9922746

Fajen, B.R., & Warren, W.H. Behavioral dynamics of steering, obstacle avoidance, and route selection. J. Exp. Psychol. Hum. Percept. Perform. 29, 343–362 (2003). 10.1037/0096-1523.29.2.343

Frohnwieser, A., Hopf, R., & Oberzaucher, E. Human walking behavior – the effect of pedestrian flow and personal space invasions on walking speed and direction. Hum. Ethol. Bull. 28, 20–28 (2013).

Gérin-Lajoie, M., Richards, C.L., & McFadyen, B.J. The negotiation of stationary and moving obstruction during walking: anticipatory locomotor adaptations and preservation of personal space. Motor. Control. 9, 242–269 (2005). 10.1123/mcj.9.3.242

Gérin-Lajoie, M., Richards, C.L., & McFadyen, B.J. The circumvention of obstacles during walking in different environmental contexts: a comparison between older and younger adults. Gait. Posture. 24, 364–369 (2006). 10.1016/j.gaitpost.2005.11.001

Gibson, J.J. The ecological approach to visual perception. (Houghton Mifflin, 1979).

Gorrini, A., Shimura, K., Bandini, S., Ohtsuka, K., & Nishinari, K. Experimental investigation of pedestrian personal space: toward modeling and simulation of pedestrian crowd dynamics. Transportation. Res. Rec. 2421, 57–63 (2014). 10.3141/2421-07

Graziano, M.S.A., & Cooke, D.F. Parieto-frontal interactions, personal space, and defensive behavior. Neuropsychologia. 44, 845–859 (2006). 10.1016/j.neuropsychologia.2005.09.009

Hackney, A.L., Cinelli, M.E., Warren, W.H., & Frank, J.S. Are avatars treated like human obstacles during aperture crossing in virtual environments? Gait. Posture. 80, 74–76 (2020). 10.1016/j.gaitpost.2020.05.028

Hartnett, J.J., Bailey, K.G., & Hartley, C.S. Body height, position, and sex as determinants of personal space. J. Psychol. 87, 129–136 (1974).

Hecht, H., Welsch, R., Viehoff, J., & Longo, M.R. The shape of personal space. Acta. Psychol. (Amst). 193, 113–122 (2019). 10.1016/j.actpsy.2018.12.009

Huber, M., Su, Y-H., Krüger, M., Faschian, K., Glasauer, S., & Hermsdörfer, J. Adjustments of Speed and Path when Avoiding Collisions with Another Pedestrian. PLoS. ONE. 9, e89589 (2014). 10.1371/journal.pone.0089589.

Kelly, J.W., Donaldson, L.S., Sjolund, L.A., & Freiberg, J.B. More than just perception—action recalibration: walking through a virtual environment causes rescaling of perceived space. Atten. Percept. Psychophys. 75, 1473–1485 (2013). 10.3758/s13414-013-0503-4

Koilias, A., Nelson, M.G., Anagnostopoulos, C.N., & Mousas, C. Immersive walking in a virtual crowd: The effects of the density, speed, and direction of a virtual crowd on human movement behavior. Comput. Animat. Virtual. Worlds. 31, e1928 (2020). 10.1002/cav.1928

Lamberg, E.M., & Muratori, L.M. Cell phones change the way we walk. Gait. Posture. 35, 688–690 (2012). 10.1016/j.gaitpost.2011.12.005

Lin, M-I., & Huang, Y-P. The impact of walking while using a smartphone on pedestrians’ awareness of roadside events. Accid. Anal. Prev. 101, 87–96 (2017). 10.1016/j.aap.2017.02.005

Liu, M.W., Wang, S.M., Oeda, Y., & Sumi, T.N. Simulating uni-and bi-directional pedestrian movement on stairs by considering specifications of personal space. Accid. Anal. Prev. 122, 350–364 (2019). 10.1016/j.aap.2017.11.012

Murakami, H., Tomaru, T., Feliciani, C., & Nishiyama, Y. Spontaneous behavioral coordination between avoiding pedestrians requires mutual anticipation rather than mutual gaze. iScience. 25, 105474 (2022). 10.1016/j.isci.2022.105474

Nikmanesh, M., Cinelli, M.E., & Marigold, D.S. Identifying factors that contribute to collision avoidance behaviours while walking in a natural environment. bioRxiv. (2024). 10.1101/2024.06.11.598509

Pfaff, L.M., & Cinelli, M.E. Avoidance behaviours of young adults during a head-on collision course with an approaching person. Exp. Brain. Res. 236, 3169–3179 (2018). 10.1007/s00221-018-5371-7

Rapos, V., Cinelli, M., Snyder, N., Crétual, A., & Olivier, A-H. Minimum predicted distance: Applying a common metric to collision avoidance strategies between children and adult walkers. Gait. Posture. 72, 16–21 (2019). 10.1016/J.GAITPOST.2019.05.016

Rapos, V., Cinelli, ME., Grunberg, R., Bourgaize, S., Crétual, A., & Olivier, A-H. Collision avoidance behaviours between older adult and young adult walkers. Gait. Posture. 88, 210–215 (2021). 10.1016/J.GAITPOST.2021.05.033

Sanz, F.A., Olivier, A-H., Bruder, G., Pettré, J., & Lécuyer, A. Virtual proxemics: locomotion in the presence of obstacles in large immersive projection environments. 2015 IEEE Virtual Reality (VR). 23-27 (2015).

Shi, Z., Zhang, J., Shang, Z., & Song, W. Collision avoidance behaviours of luggage-laden pedestrians. Physica. A. 639, 129664 (2024). 10.1016/j.physa.2024.129664

Shimizu, K., Kihara, Y., Itou, K., Tai, K., & Furuna, T. How perception of personal space influence obstacle avoidance during walking: differences between young and older adults. Phys. Ther. Res. 23, 31–38 (2020). 10.1298/ptr.E9988.

Souza Silva, W., Aravind, G., Sangani, S., & Lamontagne, A. Healthy young adults implement distinctive avoidance strategies while walking and circumventing virtual human vs. non-human obstacles in a virtual environment. Gait. Posture. 61, 294–300 (2018).

Souza Silva, W., McFadyen, B., Kehayia, E., Azevedo, N., Fung, J., & Lamontagne, A. Phone messages affect the detection of approaching pedestrians in healthy young and older adults immersed in a virtual community environment. PLoS. ONE. 14, e0217062 (2019). 10.1371/journal.pone.0217062

Souza Silva, W., McFadyen, B., Fung, J., & Lamontagne, A. Reading text messages at different stages of pedestrian circumvention affects strategies for collision avoidance in young and older adults. Gait. Posture. 76, 290–297 (2020).

Willemsen, P., Colton, M.B., Creem-Regehr, S.H., & Thompson, W.B. The effects of head-mounted display mechanical properties and field of view on distance judgments in virtual environments. ACM. Trans. Appl. Percept. 6, Article 8 (2009). http://doi.acm.org/10.1145/1498700.1498702

